# Applying AlphaGenome Variant Impact for SNV Prioritization

**DOI:** 10.64898/2026.09.26.754650

**Authors:** Hui-Qi Qu, Hakon Hakonarson

**Author notes:** Correspondence: Dr. Hakon Hakonarson Center for Applied Genomics 3615 Civic Center Blvd Abramson Building, Philadelphia, PA 19104, United States of America.

## Abstract

AlphaGenome Atlas, released in September 2026, provides functional predictions for nearly all possible human single-nucleotide variants using the AlphaGenome Variant Impact (AVI) score. However, empirical thresholds relating AVI quantile rank (AVI_QUANTILE) to clinical variant classifications have not been established. We derived AVI_QUANTILE thresholds distinguishing ClinVar pathogenic (P), likely pathogenic (LP), uncertain, conflicting, likely benign, and benign SNVs using receiver operating characteristic analyses. P/LP variants were strongly enriched at the upper end of the AVI_QUANTILE distribution, yielding a Youden-optimal ClinVar-derived cutoff of 0.9938. Optimal cutoffs varied across molecular consequence classes, with higher thresholds for missense variants and lower thresholds for intronic and UTR variants. Most well-powered gene-disease groups showed cutoffs close to the global threshold, with modest gene-specific variation. Among 24,989 variants of uncertain significance, 11,356 (45.44%) exceeded the global cutoff and had AVI_QUANTILE values within the range of P/LP variants. In a steroid-dependent asthma fine-mapping analysis, AVI_QUANTILE showed little correspondence with SuSiE posterior inclusion probabilities, indicating that AlphaGenome functional predictions capture information distinct from statistical fine-mapping.

## Introduction

Clinical variant classification integrates multiple forms of evidence, including population frequency, functional data, segregation, phenotype consistency, and computational prediction (Richards et al. 2015). Variants without sufficient or consistent evidence are classified as variants of uncertain significance (VUS), for which additional evidence is required before confident interpretation. ClinVar provides a large public archive of clinical variant interpretations and has become a valuable source for training and evaluating computational models of variant classification (Landrum et al. 2018).

Computational predictions can contribute to variant interpretation when experimental or family-based evidence is limited. ClinGen has established quantitative calibration approaches for assigning PP3/BP4 evidence from computational predictors (Pejaver et al. 2022). AlphaMissense predicts the effects of missense variants on protein function using structural and evolutionary information (Cheng et al. 2023). AlphaGenome models sequence effects on gene expression, splicing, chromatin accessibility, transcription-factor binding, histone modifications, chromatin contacts, and other regulatory features (Avsec et al. 2026). AlphaGenome is a genomic foundation model developed by Google DeepMind. The AlphaGenome Variant Impact (AVI) score combines predictions from AlphaGenome and AlphaMissense into a single variant-impact score, allowing both regulatory and protein-altering effects to contribute to variant prioritization (AlphaGenome Atlas 2026).

In September 2026, AlphaGenome Atlas released precomputed predictions for approximately 9 billion possible human single-nucleotide variants, covering nearly every possible single-base substitution in the genome (AlphaGenome Atlas 2026). The Atlas includes AVI quantile rank (AVI_QUANTILE), a standardized measure relative to the scorer reference distribution. Precomputed AVI scores permit evaluation of large variant sets without individual model inference. We evaluated AlphaGenome Atlas AVI scores across ClinVar SNVs and derived empirical AVI_QUANTILE thresholds using pathogenic/likely pathogenic (P/LP) and benign/likely benign (B/LB) classifications. We assessed VUSs using the threshold and compared AVI_QUANTILE with statistical fine-mapping results in steroid-dependent asthma.

## Methods

### AlphaGenome Atlas annotation

ClinVar data were obtained from the September 13, 2026 release. Variants were grouped according to ClinVar CLNSIG. All pathogenic and likely pathogenic SNVs were included. For comparison groups, 25,000 SNVs were randomly sampled from each of four categories: uncertain significance, conflicting classifications of pathogenicity, likely benign, and benign. Analyses were restricted to variants with A, C, G, or T as both reference and alternate alleles. Variants were annotated using AlphaGenome Atlas through the AlphaGenome Python package version 0.9.0 (Google 2026).

### Receiver operating characteristic (ROC) analysis

ROC analysis was performed to derive empirical AVI_QUANTILE thresholds for distinguishing pathogenic/likely pathogenic SNVs from benign/likely benign SNVs. Sensitivity and specificity were calculated at each cutoff, and the optimal cutoff was defined by the maximum Youden index. Thresholds corresponding to prespecified sensitivity and specificity levels were also examined. All analyses applying the global cutoff used its full numerical precision, with rounding for presentation. Across six ClinVar classes: P, LP, VUS, conflicting classifications, LB, and B, differences were assessed using the Kruskal-Wallis test followed by pairwise Mann-Whitney U tests with Holm correction. ROC analyses were further performed separately for molecular consequence categories, including missense, synonymous, intronic, UTR, stop-gained, and canonical splice variants, when at least 50 pathogenic/likely pathogenic and 50 benign/likely benign variants were available.

Gene-disease-specific analyses were performed for individual gene-disease pairs. ROC analyses were performed for gene-disease groups containing at least 100 pathogenic/likely pathogenic and 50 benign/likely benign SNVs. Performance at the global AVI_QUANTILE cutoff was also evaluated within each eligible group.

### Application to fine-mapping of steroid-dependent asthma

AVI_QUANTILE was evaluated in our steroid-dependent asthma (SDA) association analysis of the chromosome 1q31 region (Qu et al. 2026). Association analyses had been performed separately in African-ancestry (AFR) and European-ancestry (EU) samples, followed by ancestry-specific SuSiE-RSS fine-mapping using ancestry-matched linkage disequilibrium (Wang et al. 2020; Zou et al. 2022). Seven non-overlapping regions were fine-mapped in each ancestry, with 95% credible sets generated within each region. SuSiE posterior inclusion probability (PIP), credible-set membership, and AVI_QUANTILE were evaluated jointly.

Spearman correlation was calculated between PIP and AVI_QUANTILE across all annotated variants. AVI_QUANTILE distributions and the proportion of variants exceeding the ClinVar-derived threshold of 0.9938 were compared between credible-set and non-credible-set variants. Within multi-variant credible sets, the variant with the highest PIP was compared with the variant with the highest AVI_QUANTILE. Significance of the observed number of matches was evaluated using the exact Poisson-binomial distribution and Monte Carlo simulation using the corresponding set-specific probabilities.

## Results

### AVI_QUANTILE between ClinVar P/LP and B/LB

A total of 84,730 P, 78,468 LP, 24,989 VUS, 24,997 variants with conflicting classifications, 24,984 LB, and 24,879 B variants were included after AlphaGenome annotation, respectively (Table 1). P and LP variants were concentrated at the upper end of the AVI_QUANTILE distribution. Median AVI_QUANTILE values were 0.9995 for P and 0.9990 for LP, compared with 0.8198 for LB and 0.6588 for B. At the more stringent threshold of 0.9990, 66.61% of P and 51.27% of LP variants exceeded the threshold, compared with 0.07% of LB and 0.17% of B variants.

**Table 1.** AVI_QUANTILE summary across ClinVar classes.

| Class | N | Median | Q1 | Q3 | ≥0.90 (%) | ≥0.95 (%) | ≥0.99 (%) | ≥0.999 (%) |
|---|---|---|---|---|---|---|---|---|
| **Pathogenic** | 84,730 | 0.99952 | 0.99856 | 0.99981 | 99.46 | 99.22 | 97.74 | 66.61 |
| **Likely pathogenic** | 78,468 | 0.99904 | 0.99798 | 0.99956 | 99.49 | 99.22 | 97.57 | 51.27 |
| **VUS** | 24,989 | 0.99268 | 0.97176 | 0.99631 | 85.94 | 80.68 | 58.43 | 4.00 |
| **Conflicting** | 24,997 | 0.98260 | 0.85691 | 0.99523 | 71.16 | 63.60 | 41.62 | 4.66 |
| **Likely benign** | 24,984 | 0.81982 | 0.24872 | 0.95513 | 40.62 | 27.11 | 5.24 | 0.07 |
| **Benign** | 24,879 | 0.65880 | 0.16273 | 0.93416 | 30.44 | 21.86 | 7.62 | 0.17 |

### Empirical AVI_QUANTILE threshold for P/LP SNVs

ROC analysis was performed using P/LP variants as the positive class and B/LB variants as the negative class. The ROC AUC was 0.9914 (Table 2; Figure 1). The optimal threshold based on the Youden index was an AVI_QUANTILE of 0.9938. At this cutoff, sensitivity was 96.06% and specificity was 96.65%, corresponding to a Youden index of 0.9270. Of 163,198 P/LP variants, 156,764 exceeded the threshold, compared with 1,672 of 49,863 B/LB variants.

**Table 2.** ROC-derived AVI_QUANTILE thresholds for P/LP vs B/LB.

| Criterion | Cutoff | Sensitivity | Specificity |
|---|---|---|---|
| **Youden optimal** | 0.99379 | 96.06% | 96.65% |
| **90% sensitivity** | 0.99670 | 90.00% | 99.16% |
| **95% sensitivity** | 0.99484 | 95.00% | 97.63% |
| **99% sensitivity** | 0.96674 | 99.00% | 81.42% |
| **90% specificity** | 0.98403 | 98.37% | 90.01% |
| **95% specificity** | 0.99171 | 97.14% | 95.00% |
| **99% specificity** | 0.99646 | 91.13% | 99.00% |

\* Overall ROC AUC = **0.9914.**

**Figure 1.**
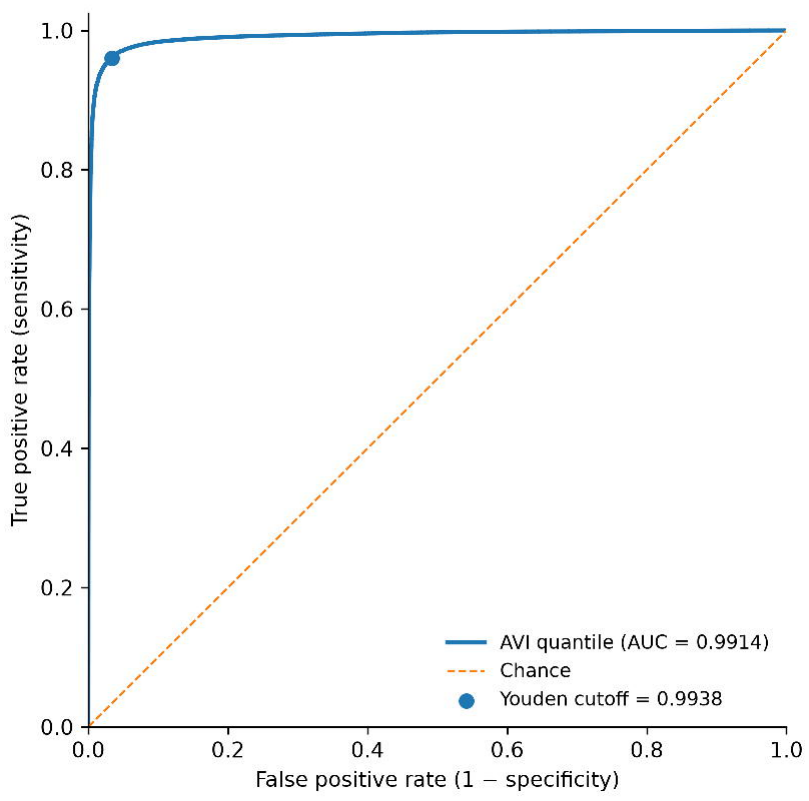
Receiver operating characteristic analysis of AVI quantile for distinguishing pathogenic/likely pathogenic from benign/likely benign SNVs. The receiver operating characteristic curve was generated using AVI_QUANTILE as the predictor. The area under the curve was 0.9914. The optimal cutoff based on the Youden index was 0.9938, corresponding to 96.1% sensitivity and 96.6% specificity.

An AVI_QUANTILE cutoff of 0.9948 provided 95.00% sensitivity and 97.63% specificity, and a cutoff of 0.9965 provided 99.00% specificity with 91.13% sensitivity. A lower cutoff of 0.9667 achieved 99.00% sensitivity with 81.42% specificity. At fixed thresholds, AVI_QUANTILE ≥0.9900 yielded 97.66% sensitivity and 93.57% specificity, AVI_QUANTILE ≥0.9950 yielded 94.78% sensitivity and 97.76% specificity, and AVI_QUANTILE ≥0.9990 yielded 59.24% sensitivity and 99.88% specificity. Because ClinVar has been used in the performance evaluation of AlphaGenome and AVI, these performance estimates may be inflated by data reuse. They are useful for defining a prioritization threshold, but not for estimating prospective clinical performance in an independent dataset.

### AVI distribution across VUS and conflicting classifications

VUS and variants with conflicting classifications showed intermediate AVI_QUANTILE distributions (Table 1). Median values were 0.9927 for VUS and 0.9826 for variants with conflicting classifications. Using the cutoff of 0.9938, 11,356 of 24,989 VUSs (45.44%) and 8,078 of 24,997 variants with conflicting classifications (32.32%) exceeded the threshold.

### AVI performance across molecular consequence categories

For missense variants, the optimal cutoff was 0.9957, with an AUC of 0.9541, sensitivity of 88.97%, and specificity of 90.07%. Synonymous variants had an optimal cutoff of 0.9898 (AUC = 0.9006; sensitivity = 73.02%; specificity = 96.52%), and intronic variants had an optimal cutoff of 0.9744 (AUC = 0.9768; sensitivity = 91.39%; specificity = 96.04%). UTR variants had a lower optimal cutoff of 0.9536, with an AUC of 0.8463, sensitivity of 72.09%, and specificity of 85.96% (Table 3).

**Table 3.**
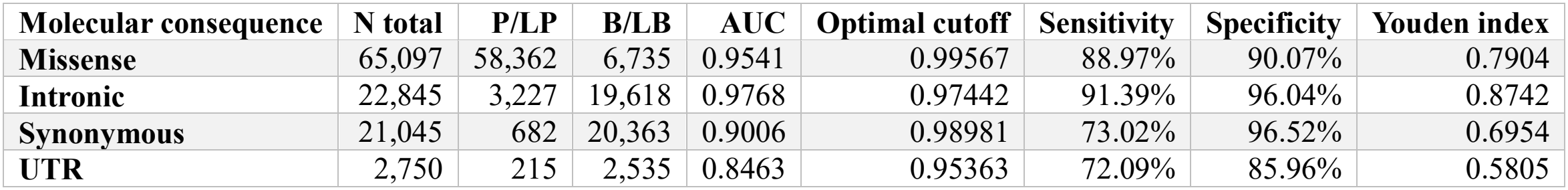
AVI_QUANTILE performance across molecular consequence categories.

| **Molecular consequence** | **N total** | **P/LP** | **B/LB** | **AUC** | **Optimal cutoff** | **Sensitivity** | **Specificity** | **Youden index** |
|---|---|---|---|---|---|---|---|---|
| **Missense** | 65,097 | 58,362 | 6,735 | 0.9541 | 0.99567 | 88.97% | 90.07% | 0.7904 |
| **Intronic** | 22,845 | 3,227 | 19,618 | 0.9768 | 0.97442 | 91.39% | 96.04% | 0.8742 |
| **Synonymous** | 21,045 | 682 | 20,363 | 0.9006 | 0.98981 | 73.02% | 96.52% | 0.6954 |
| **UTR** | 2,750 | 215 | 2,535 | 0.8463 | 0.95363 | 72.09% | 85.96% | 0.5805 |

Reliable ROC-based cutoff estimation was not possible for stop-gained and canonical splice variants because ClinVar contained very few B/LB variants in these categories. Among P/LP variants, 97.32% of stop-gained variants and 99.47% of canonical splice variants met or exceeded an AVI_QUANTILE cutoff of 0.994.

### Performance across gene-disease groups

Gene-disease-specific analyses were performed for 13 gene-disease groups (Table 4). AUC values ranged from 0.9706 for *DNAH11*-associated primary ciliary dyskinesia to 1.0000 for a *TTN*-associated disease group. Optimal thresholds ranged from 0.9908 for *TSC2* to 0.9977 for *DMD*. At the global cutoff, sensitivity ranged from 90.69% to 100% and specificity from 86.01% to 100%.

**Table 4.** Performance of AVI_QUANTILE across gene-disease groups.

| **Gene-disease group\*** | **P/LP** | **B/LB** | **AUC** | **Optimal cutoff** | **Sens.** | **Spec.** | **Sens. at global 0.99379** | **Spec. at global 0.99379** |
|---|---|---|---|---|---|---|---|---|
| ***TTN* - DCM/LGMD2J** | 1,203 | 234 | 0.9999 | 0.99402 | 99.75% | 100% | 99.75% | 100% |
| ***DMD* - Duchenne MD** | 648 | 59 | 0.9988 | 0.99774 | 97.84% | 100% | 99.38% | 96.61% |
| ***NEB* - Nemaline myopathy 2** | 479 | 129 | 0.9998 | 0.99629 | 99.58% | 99.22% | 100% | 95.35% |
| ***VPS13B* - Cohen syndrome** | 355 | 67 | 0.9981 | 0.99619 | 97.75% | 100% | 99.15% | 98.51% |
| ***DNAH11* - Primary ciliary dyskinesia** | 216 | 143 | 0.9706 | 0.99636 | 92.13% | 95.10% | 94.44% | 86.01% |
| ***PKHD1* - AR polycystic kidney disease** | 303 | 52 | 0.997 | 0.99216 | 98.68% | 100% | 98.02% | 100% |
| ***DNAH5* - Primary ciliary dyskinesia** | 277 | 61 | 0.9987 | 0.99647 | 98.19% | 98.36% | 99.28% | 95.08% |
| ***APC* - Familial adenomatous polyposis 1** | 262 | 63 | 0.9995 | 0.9964 | 99.24% | 100% | 99.24% | 93.65% |
| ***TSC2* - Tuberous sclerosis 2** | 204 | 67 | 0.9809 | 0.99083 | 93.63% | 94.03% | 90.69% | 94.03% |
| ***TTN* - DCM/LGMD2J + cardiovascular phenotype** | 135 | 73 | 1 | 0.99616 | 100% | 100% | 100% | 100% |
| ***BRCA2* - HBOC susceptibility 2** | 134 | 62 | 0.993 | 0.99324 | 96.27% | 100% | 94.78% | 100% |
| ***LYST* - Chédiak-Higashi syndrome** | 122 | 56 | 0.993 | 0.99668 | 99.18% | 100% | 99.18% | 98.21% |
| ***BRCA1* - HBOC susceptibility 1** | 112 | 56 | 0.9997 | 0.99622 | 98.21% | 100% | 98.21% | 98.21% |

\* Only gene-disease groups containing at least 100 P/LP and 50 B/LB SNVs were included. P/LP, pathogenic/likely pathogenic; B/LB, benign/likely benign; DCM, dilated cardiomyopathy; LGMD2J, limb-girdle muscular dystrophy type 2J; HBOC, hereditary breast and ovarian cancer; AR, autosomal recessive.

### Nonredundant information from AlphaGenome and statistical fine-mapping

A total of 41,156 ancestry-specific SuSiE-analyzed SNV records were annotated with AlphaGenome, including 23,152 AFR and 18,004 EU records representing 30,539 distinct SNVs. Of these, 273 belonged to 84 SuSiE 95% credible sets (Table 5). AVI_QUANTILE showed little correlation with SuSiE PIP. The overall Spearman correlation was 0.0434, with similarly small correlations in AFR (ρ = 0.0417, P = 2.21 × 10^-10^) and EU (ρ = 0.0436, P = 4.96 × 10^-9^).

**Table 5.** Comparison of AlphaGenome AVI_QUANTILE with SuSiE fine-mapping results in steroid-dependent asthma.

| **Analysis** | **Overall** | **AFR** | **EU** |
|---|---|---|---|
| **SuSiE-analyzed SNVs with AVI** | 41,156 | 23,152 | 18,004 |
| **Credible-set SNVs** | 273 | 183 | 90 |
| **Spearman rho, PIP vs AVI** | 0.0434 | 0.0417 | 0.0436 |
| **Median AVI, CS** | 0.4676 | 0.4676 | 0.4659 |
| **Median AVI, non-CS** | 0.393 | 0.4035 | 0.3788 |
| **CS vs non-CS P** | 0.0219 | 0.1294 | 0.0902 |
| **AVI ≥0.9938 in CS** | 0/273 | 0/183 | 0/90 |
| **Multi-variant CS** | 46 | 25 | 21 |
| **Top-PIP = top-AVI** | 15/46 (32.6%) | 7/25 (28.0%) | 8/21 (38.1%) |
| **Expected by chance** | 31.40% | 26.40% | 37.40% |
| **Exact P** | 0.8668 | 1 | 1 |

The median AVI_QUANTILE was 0.4676 among credible-set variants and 0.3930 among non-credible-set variants (Mann–Whitney P = 0.0219). Differences were not significant when evaluated separately in AFR (P = 0.1294) or EU (P = 0.0902). None of the 273 credible-set variants met the ClinVar-derived AVI_QUANTILE threshold of 0.9938, compared with 55 of 40,883 non-credible-set variants (0.13%; Fisher’s exact P = 1.000).

Among 84 credible sets, 38 contained a single SNV and were uninformative for ranking agreement. Among the 46 multi-variant credible sets, the highest-PIP and highest-AVI variant was the same in 15 sets (32.6%). Based on the observed credible-set size distribution, 14.45 matches (31.4%) were expected by chance. The observed concordance did not differ from this expectation (exact P = 0.8668; Monte Carlo P = 0.8666). Concordance was not significant when evaluated separately in AFR or EU credible sets (Table 5).

## Discussion

### High AVI_QUANTILE in P/LP variants

AVI integrates regulatory and transcript-level predictions from AlphaGenome with protein-effect predictions from AlphaMissense into a common impact score (Cheng et al. 2023; AlphaGenome Atlas 2026; Avsec et al. 2026). High AVI_QUANTILE values were observed across pathogenic coding and noncoding variant classes. AVI_QUANTILE should not be interpreted as a probability of pathogenicity. It is a rank relative to the AVI reference distribution, so a value near 0.994 places a variant near the upper extreme of predicted molecular impact without implying a corresponding probability of disease causation. Clinical classification requires integration of computational prediction with population frequency, phenotype, segregation, functional evidence, and other lines of evidence (Richards et al. 2015; Pejaver et al. 2022). The ClinVar-derived cutoff can be used as a variant-prioritization threshold rather than a standalone classification criterion. Its high apparent discrimination may partly reflect overlap between information represented in ClinVar and information used during model development or calibration, and prospective performance will need to be established in fully independent datasets.

### Variation across molecular consequence classes and gene-disease groups

Optimal thresholds varied across molecular consequence classes, with lower thresholds for intronic and UTR variants. Intronic variants showed the highest AUC, followed by missense variants, with lower AUCs for synonymous and UTR variants. These findings support molecular consequence-specific calibration of AVI scores.

Variation among gene-disease groups may similarly reflect differences in underlying disease mechanisms, gene constraint, and variant composition. The relative importance of missense, truncating, splice, and other variant classes differs among genes and disorders. ClinGen expert panels already use gene-specific adaptations of variant interpretation criteria (Richardson et al. 2024). Our findings support gene- and mechanism-specific calibration rather than assuming that a single global threshold will perform equivalently across all clinical contexts. However, robust estimation of such thresholds will require sufficiently large independent datasets.

### High AVI_QUANTILE in a subset of VUSs

VUSs are inherently heterogeneous in predicted molecular effect, population frequency, segregation evidence, phenotype consistency, and available functional data. Strong predicted molecular effects may help prioritize VUSs for functional assays, segregation analysis, or phenotype reassessment. ClinGen recommendations allow appropriately calibrated computational evidence to contribute to PP3/BP4, but not to independently determine pathogenicity (Pejaver et al. 2022). Current ACMG guidance emphasizes that additional evidence is required before a VUS can be assigned a definitive clinical interpretation (Mighton et al. 2026).

Among VUSs, 45.44% exceeded the ClinVar-derived threshold and occupied a region of the AVI_QUANTILE distribution enriched for P/LP variants. These variants are not necessarily pathogenic, but they represent a subset that may warrant closer review using additional clinical, segregation, or functional evidence.

### AlphaGenome and statistical fine-mapping

The SDA analysis showed little correspondence between statistical evidence for association and predicted functional effect. SuSiE PIP reflects the posterior probability assigned to a variant as a causal contributor to an association signal under the fine-mapping model and local linkage disequilibrium (Wang et al. 2020; Zou et al. 2022), whereas AVI_QUANTILE reflects predicted biological and molecular effects. The weak correspondence between SuSiE PIP and AVI reflects their different objectives: statistical fine-mapping asks which variants are most compatible with the association data, while AlphaGenome predicts the magnitude of variant-associated molecular effects. Genotypes in our study were imputed using the TOPMed reference panel, which provides improved coverage of low-frequency and rare variants in African-ancestry populations, although very rare or poorly represented variants may still be missed (Taliun et al. 2021). Many common disease-associated variants have modest regulatory effects that depend on tissue, cell type, developmental stage, or other biological context (D’Antonio et al. 2023). Such context dependence may explain why none of the credible-set variants reached the ClinVar-derived AVI_QUANTILE threshold despite higher median AVI_QUANTILE values within credible sets. Complex-trait variants may have smaller molecular effects than highly penetrant Mendelian P/LP variants. A variant with a strong predicted molecular effect need not have the highest statistical support within an association signal, particularly when correlated variants tag one or more functional alleles in linkage disequilibrium (Long et al. 2025). A variant with strong statistical support may exert a subtle, context-dependent regulatory effect and need not receive an extreme AVI_QUANTILE score.

These observations suggest that AlphaGenome and statistical fine-mapping should be viewed as complementary. Combining statistical evidence of association with predicted molecular consequence may provide greater resolution for prioritizing causal variants than either approach alone.

## Limitations

ClinVar has been used in the performance evaluation of AlphaGenome and AVI, so the present analysis is not an independent external validation. Potential information leakage through benchmark reuse may lead to overestimation of discrimination. ClinVar classifications may themselves incorporate computational evidence, creating additional dependence between predictor scores and reference classifications. The AVI_QUANTILE thresholds reported here require evaluation in independent variant datasets not used in model development or threshold derivation.

## Data Availability

ClinVar data used in this study are publicly available from the NCBI ClinVar database. AlphaGenome Atlas was queried using the AlphaGenome Python package (version 0.9.0). Data from our steroid-dependent asthma analysis are described in the referenced study.

